# Pathogenicity determinant protein E is a *Francisella* type VI secretion system effector protein that modulates host death

**DOI:** 10.1101/2024.11.25.624902

**Authors:** Nicole Adams, Danielle Hulvey, Nimra Khalid, Alexandra Burne, Mary B. Brown, Aria Eshraghi

**Affiliations:** Department of Infectious Diseases & Immunology, University of Florida, Gainesville, Florida, USA; Emerging Pathogens Institute, University of Florida, Gainesville, Florida, USA; Department of Oral Biology, University of Florida, Gainesville, Florida, USA

## Abstract

*Francisella tularensis* is a highly pathogenic Gram-negative intracellular bacterial pathogen and the etiologic agent of tularemia, a fatal zoonotic disease that poses a threat to global public health. *F. tularensis* virulence is mediated by genes on the *Francisella* pathogenicity island (FPI), which encodes a unique contractile secretion apparatus with functional and structural similarity to bacterial type VI secretion systems (T6SSs). T6SSs can inject effector proteins into host cells to facilitate invasion, intracellular proliferation, and pathogenesis; however, the identity and function of *Francisella* effectors are still unclear. In this study, we measured T6SS activity in a series of FPI deletion mutants to identify genes that are required for core apparatus activity. We found that Pathogenicity Determinant Protein E (PdpE) is not required for secretion of T6SS substrates or intramacrophage growth. Instead, PdpE forms a complex with another T6SS-secreted effector, PdpC, and limits death of infected host cells by attenuating the macrophage type I interferon response. Importantly, a Δ*pdpE* mutant more rapidly induces death in an established invertebrate *in vivo* model for *Francisella* infection. These data define the role of a previously uncharacterized substrate of the *Francisella* T6SS and provide new insights into host-*Francisella* interaction.

## Introduction

*Francisella* are host-adapted Gram-negative intracellular bacterial pathogens that cause septicemia in a broad range of hosts, including humans. *F. tularensis* subspecies *tularensis* (*F. tularensis*) are transmitted through multiple routes and exhibit a remarkable virulence that is exemplified by a lethal dose below 10 viable bacteria and greater than 60% mortality if left untreated. *F. tularensis* virulence is mediated by replication within macrophages and requires genes on the *Francisella* pathogenicity island (FPI), which encodes an intercellular protein delivery apparatus that is structurally and functionally related to the bacterial type VI secretion system (T6SS) [1]. While T6SSs are widespread in proteobacteria and are predominantly associated with interbacterial competition, some target both bacteria and host cells. The FPI-encoded T6SS is phylogenetically distinct, unique to *Francisella*, and exclusively targets host cells to promote virulence; thus, it has been classified as a separate subtype, the T6SS^ii^ [2].

Host-targeted T6SSs are composed of three classes of proteins: 1) structural components of the apparatus required for T6SS activity, 2) secreted proteins required for T6SS activity, and 3) T6SS effector proteins, which are secreted into host cells to induce physiological changes that benefit the pathogen. By studying the extracellular proteome of a closely related subspecies that encodes a similar FPI to *F. tularensis, F. novicida*, we identified T6SS-exported proteins and differentiated between those required for T6SS activity (class 2) and others that are not required for apparatus activity and may be effectors (class 3) [3]. Loss-of-function mutations in effector genes does not affect core apparatus activity. However, deletion of all known effector genes in a single strain renders it unable to grow within cultured macrophages, suggesting that the known effectors comprise the full complement required for pathogenicity. Based on these data, we surmised that the remaining genes in the FPI encode structural components of the T6SS apparatus. Here we utilized a systematic approach to define the contribution of individual FPI genes to apparatus activity, which led to the unexpected finding that Pathogenicity Determinant Protein E (PdpE), a protein with previously unknown function, interacts with a key effector, PdpC, and is secreted in a T6SS-dependent manner. Furthermore, we discovered that a Δ*pdpE* mutant induces greater cytotoxicity *in vitro* and pathogenicity in an *in vivo* model of infection. Taken together, these data reveal a new role for a previously uncharacterized FPI-encoded gene that impacts *Francisella* host-pathogen interactions.

## Results and Discussion

### Identification of FPI-encoded genes required for core T6SS activity

Proteins encoded near T6SS apparatus genes are often secreted effectors; therefore, genes within the FPI that are dispensable for core T6SS activity represent candidate effector genes. In our search for FPI-encoded effectors, we engineered *F. novicida* mutants with in-frame chromosomal deletions of each FPI gene and measured secretion of a well-characterized T6SS substrate, IglC, into the supernatant (Figure 1A). Deletion of the genes at the 5’ ends of both the upstream and downstream operons in the FPI (FTN_1309-FTN_1318 and FTN_1321-FTN_1324, respectively) was sufficient to inhibit IglC secretion; indicating that these genes are required for core apparatus function (Figure 1B). Conversely, IglC secretion was unaffected by deletion of the remaining genes, *pdpC* (FTN_1319), *pdpE* (FTN_1320), *pdpD* (FTN_1325), and *anmK* (FTN_1326). We previously determined that *pdpC, pdpD*, and two genes outside the FPI, *opiA* and *opiB*, encode effectors and are not required for T6SS activity [3]. Deletion of *pdpE* does not attenuate IglC secretion; therefore, it may be a secreted effector that has yet to be characterized.

**Figure 1:**
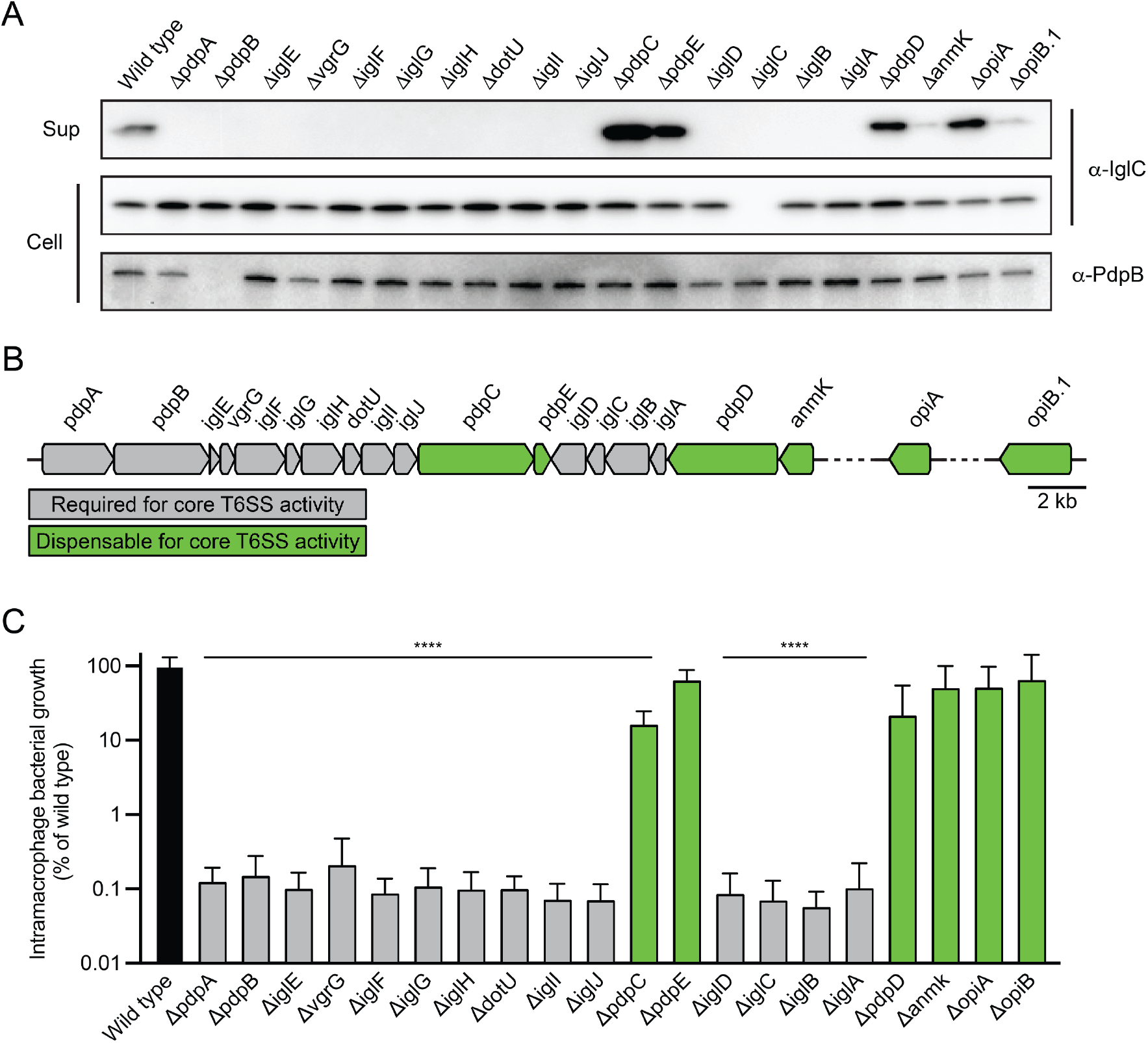
PdpE is not required for T6SS apparatus activity and is dispensable for intramacrophage growth. **(A)** Core function of the T6SS apparatus measured by western blot analysis of supernatant fractions (sup) and cell pellets from the indicated *F. novicida* strains. Western blot was performed with antibodies recognizing a secreted T6SS substrate, IglC, and membrane-associated apparatus protein, PdpB. Data are representative of four biological replicates. **(B)** Regions of the *F. novicida* genome that encode T6SS-related genes, including the *Francisella* pathogenicity island (FTN_1309 to FTN_1326) and two additional loci outside of the pathogenicity island (FTN_0131 and FTN_1069). Genes that are required or dispensable for core T6SS activity are colored grey and green, respectively. **(C)** Growth of the indicated *F. novicida* strains within immortalized murine bone marrow-derived macrophages after 18 hours. These data represent four biological replicates, each of which are composed of at least four technical replicates. (Student’s t-test; ^****^ P < 0.0001)

*F. novicida* requires the T6SS for intracellular growth; thus, we quantified bacterial replication within immortalized bone marrow-derived macrophages (iBMDMs) as an *in vivo* measurement of the contribution of FPI genes to T6SS activity. Deletion of genes required for IglC secretion severely attenuates the growth of these strains in iBMDMs (Figure 1C). In contrast, deletion of *pdpE* has little to no effect on intracellular growth, similar to known effectors. These results are consistent with PdpE having the role of an effector because it is dispensable for apparatus function and phenocopies other effectors.

### PdpE interacts with PdpC and is secreted via the T6SS

*Francisella* species belong to two phylogenetically distinct groups that occupy terrestrial and marine habitats, and without exception, the strains that encode *pdpE* are limited to the terrestrial clade [4]. *pdpE* is closely linked to *pdpC*, as they are encoded adjacent to one another in all sequenced genomes except *F. opportunistica*, a strain with unclear virulence potential that lacks a functional *pdpC* gene (Figure 2A) [5,6]. Based on these data, we surmised that PdpC and PdpE may interact to form a complex. Indeed, we readily detected PdpE in association with PdpC by immunoprecipitation followed by mass spectrometry (Figure 2B). To validate these results, we performed a reciprocal pulldown of polyhistidine-tagged PdpE on Ni-NTA beads followed by western blot, confirming that PdpC co-precipitates with PdpE (Figure 2C). These data suggest that PdpE forms a secreted effector complex with PdpC.

**Figure 2:**
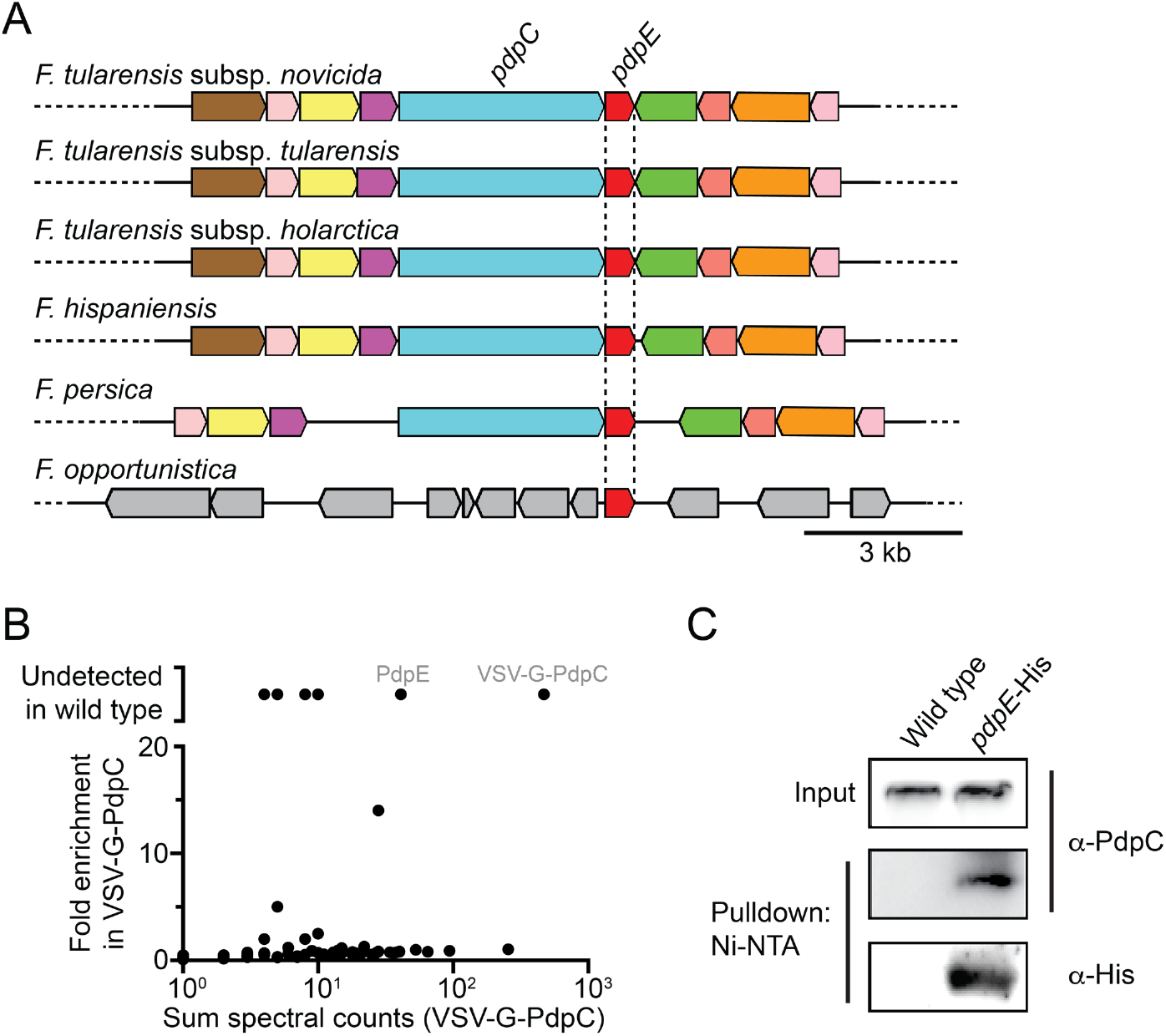
PdpE is associated with PdpC. **(A)** Genome neighborhood diagram of the loci that encode *pdpE* in multiple strains of *Francisella* generated using the Enzyme Function Initiative Genome Neighborhood Tool [22]. Homologs share colors across the strains and those colored grey are absent in these loci in other strains. **(B)** Mass spectrometry analysis and comparison of proteins that immunoprecipitate on α-VSV-G agarose beads from wild-type *F. novicida* and a VSV-G-PdpC mutant. **(C)** Western blot analysis of Ni-NTA pulldown from lysates harvested from wild-type *F. novicida* and a *pdpE*-His mutant.

Since secreted effector proteins play a major role in host-bacterial interactions and are important for virulence, their identification is a requisite step toward a mechanistic understanding of pathogenesis. Thus far, it has been unclear if PdpE is a T6SS substrate as others have reported that PdpE is secreted, not secreted, or a membrane protein [7–9]. We previously compared the extracellular proteomes of wild-type *F. novicida* with a T6SS-deficient strain (Δ*dotU*) to identify T6SS substrates [3]. In that analysis, PdpE was secreted in such low abundance that it was near our limit of detection and below our cutoff value for significance. To directly and sensitively query if PdpE is secreted via the T6SS, we inserted a VSV-G epitope DNA sequence just before the *pdpE* stop codon in wild-type *F. novicida* and Δ*dotU*, grew these strains in T6SS-activating conditions, and concentrated the supernatant fractions prior to detection of PdpE via western blot. This assay revealed that PdpE is secreted into the supernatant in wild-type *F. novicida* but not in a Δ*dotU* mutant, suggesting that PdpE is secreted via the T6SS pathway (Figure 3A).

**Figure 3:**
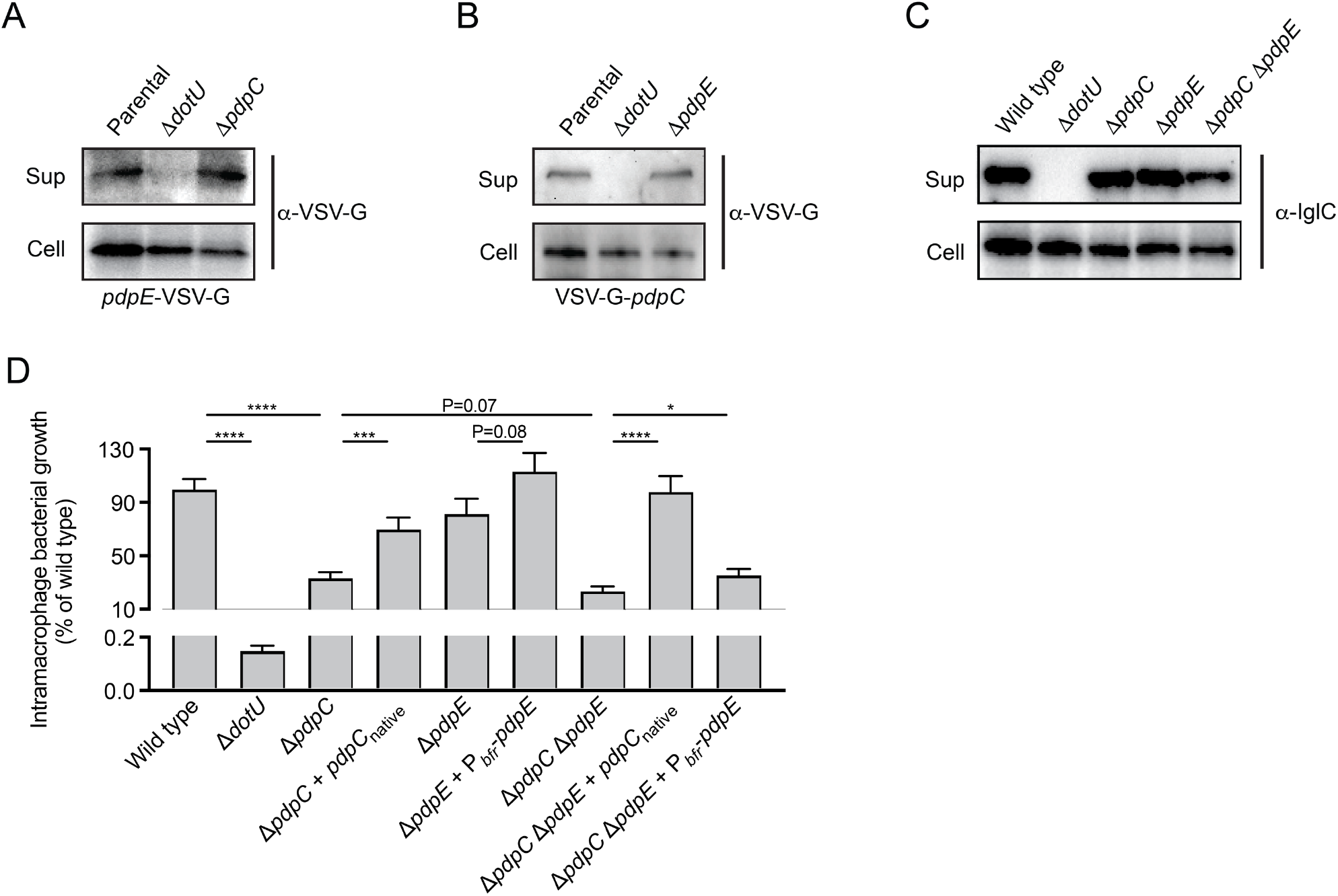
PdpE secretion is dependent on the T6SS, and expression enhances bacterial growth. **(A and B)** Western blot analysis of supernatant (sup) and cell pellets from *F. novicida* strains encoding in-frame chromosomal fusions of *pdpE* (A) and *pdpC* (B) with vesicular stomatitis virus-glycoprotein (VSV-G) at their native genomic loci. **(C)** Core function of the T6SS apparatus measured by western blot analysis of IglC in the supernatant fractions (sup) and cell pellets from the indicated genotypes. Data are representative of three biological replicates. **(D)** Growth of the indicated *F. novicida* strains within immortalized murine bone marrow-derived macrophages after 18 hours. Data represent six biological replicates, each of which are composed of four technical replicates. (Student’s t-test; ^****^ P < 0.0001, ^***^ P < 0.001, and ^*^ P < 0.05)

T6SS effector genes are often encoded adjacent to chaperones that facilitate substrate export [10,11]. Since *pdpE* is closely linked to *pdpC*–a well-known T6SS effector–we asked if they are co-dependent for secretion. Deletion of *pdpC* does not affect PdpE secretion, nor deletion of both genes (Δ*pdpC*Δ*pdpE*) does not affect apparatus activity (Figure 3C). Based on these data, we conclude that secretion of both PdpE and PdpC depend on the T6SS, but these proteins are neither co-dependent for secretion nor required for core T6SS apparatus activity.

Bacterial effectors often serve redundant functions, and in this situation only strains with multiple effector gene deletions display attenuated pathogenicity [12,13]. Indeed, deletion of *opiA* and *pdpD* only result in intracellular growth defects in Δ*pdpC* background [3,14]. Since growth of the Δ*pdpE* strain is not attenuated in macrophages, we investigated if deletion of *pdpE* has a phenotype in a Δ*pdpC* background. We validated that intracellular replication of *F. novicida* is abolished when the T6SS is genetically inactivated (*ΔdotU*) and severely attenuated by loss of *pdpC*, which is readily complemented by re-introduction of the gene into the native chromosomal locus (Figure 3D) [3,14]. Deletion of *pdpE* has no effect on intramacrophage growth; however, *pdpE* expression from a constitutive promoter modestly enhances growth, though not to a level of statistical significance. The replication defect in Δ*pdpC* is unaffected by additional loss of *pdpE* and is genetically complemented by *pdpC* expression from its native locus. Unexpectedly, expression of *pdpE* in Δ*pdpC*Δ*pdpE* modestly rescues growth, suggesting that PdpC is dispensable when PdpE is overexpressed. These results indicate that PdpE contributes to intracellular growth similar to PdpC, but likely through a different mechanism.

### PdpE modulates host death in cultured macrophages and in vivo

Since the Δ*pdpE* mutant is not attenuated for intracellular replication, we postulated that PdpE may modulate host cell death, another common role of bacterial effector proteins [13,15]. To investigate this hypothesis, we measured cytotoxicity of infected macrophages and found that Δ*pdpE* induces significantly more host cell death than wild-type *F. novicida* (Figure 4A). This phenotype is complemented by expressing *pdpE* in a heterologous site driven by either a constitutive promoter (P_bfr_) or the native FPI promoter (P_pdpA_). As expected, the T6SS-deficient strain (Δ*dotU*) does not affect host cell viability. These data suggest that PdpE inhibits cytotoxicity triggered by *F. novicida*.

**Figure 4:**
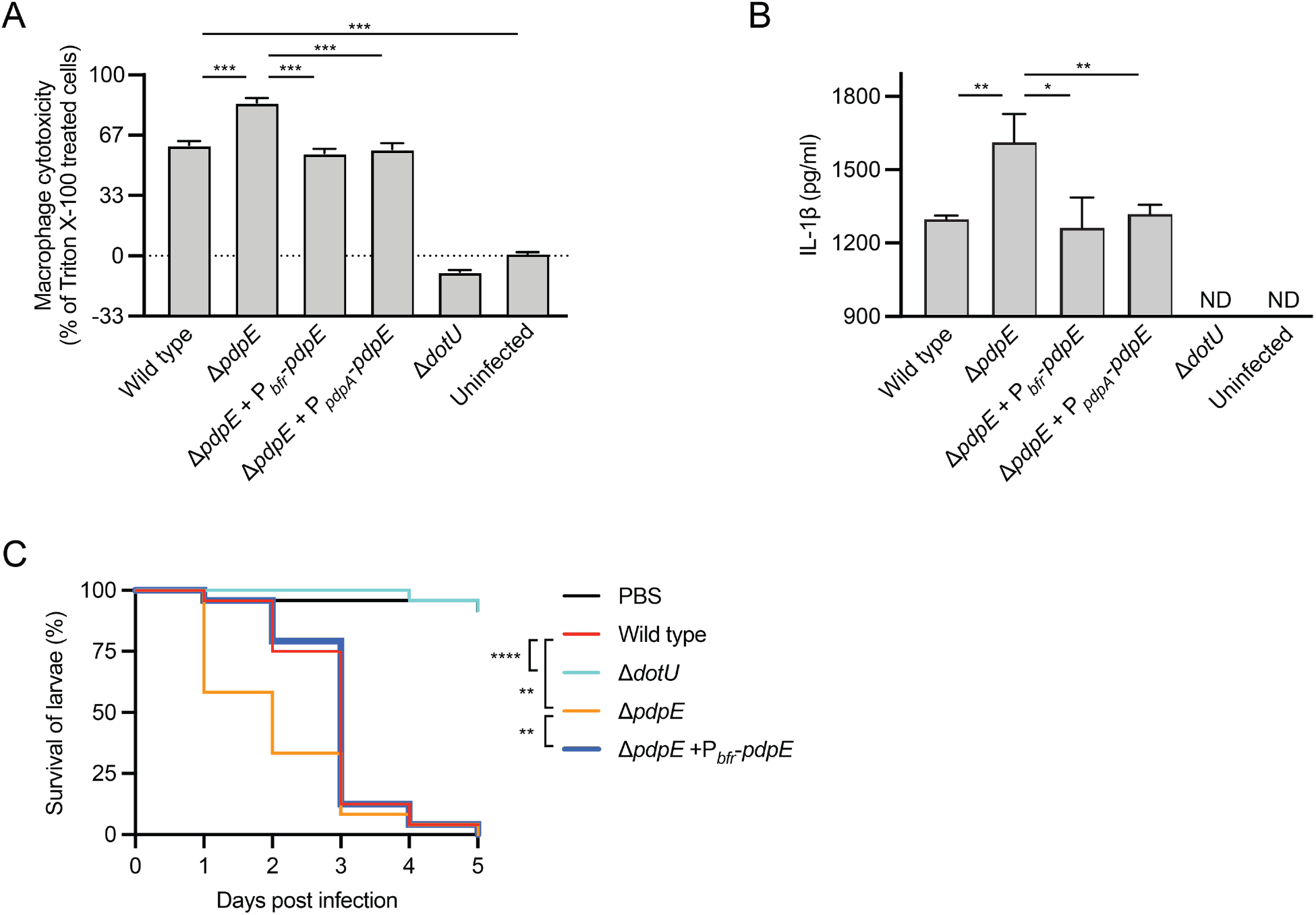
PdpE limits host death. **(A and B)** Cytotoxicity of immortalized murine bone marrow-derived macrophages (iBMDMs) as measured by release of lactate dehydrogenase into the culture supernatant 18 hours after infection with the indicated strains of *F. novicida*. **(B)** IL-1β secretion by iBMDMs pre-primed with 100 ng/mL LPS for three hours and then infected with the indicated strains of *F. novicida* for 18 hours. The data in (A) and (B) represent three biological replicates, each of which are composed of four to eight technical replicates. ND = not detected. **(C)** Survival curves of *Galleria mellonella* larvae injected with 10^4^ CFU of the indicated *F. novicida strains* and monitored two times daily. Data are representative of three biological replicates each containing at least 24 larvae. (Mantel-Cox test; ^****^ P < 0.0001 and ^**^ P < 0.002)

Intracellular *F. novicida* are recognized in the macrophage cytosol by pattern recognition receptors, which leads to activation of caspase-1, processing and secretion of IL-1β, and pyroptosis. Thus, we asked if PdpE-deficient *F. novicida* causes increased IL-1β secretion. Indeed, significantly more IL-1β was present in the supernatants of iBMDMs infected with the Δ*pdpE* strain compared to wild type (Figure 4B). Expression of *pdpE* from a constitutive promoter or the native FPI promoter (P_pdpA_) complemented IL-1β secretion, and as expected, IL-1β was not detected in the supernatants of naive iBMDMs or those infected with Δ*dotU*. These data indicate that PdpE suppresses inflammasome activity in *F. novicida*-infected macrophages, which preserves host cell viability and limits pro-inflammatory cytokine secretion.

The larval stage of *Galleria mellonella* has been used extensively as an invertebrate model for bacterial infection, including defining the importance of the T6SS pathway for *Francisella* virulence [16,17]. To test if PdpE plays a role in pathogenesis of an *in vivo* model, we compared the survival of *G. mellonella* larvae infected with wild-type *F. novicida* to a Δ*pdpE* mutant (Figure 4C). Larvae infected with wild-type *F. novicida* die rapidly, with nearly complete death within 3 days. Infection with Δ*dotU* causes almost no death, much like the larvae injected with PBS. In contrast, deletion of *pdpE* increased virulence, denoted by greater death than the wild type-infected larvae in the first 2 days post-infection. When *pdpE* was expressed in the Δ*pdpE* mutant, larval survival was nearly identical to that of wild type. These findings are consistent with our *in vitro* data and further support the role of PdpE in delaying host death.

Here we demonstrate that PdpE is a secreted effector of the *Francisella* T6SS that modulates host death both *in vitro* and *in vivo*. PdpE is secreted via the T6SS pathway and interacts with a well-characterized T6SS effector, PdpC. Consistent with being secreted T6SS effectors, both PdpE and PdpC are dispensable for core T6SS activity, but impact replication within cultured macrophages. PdpE limits and delays death of infected macrophages and *G. mellonella*. Future research will investigate the molecular mechanisms by which PdpE exerts these effects and whether it plays a role in a mammalian model for *Francisella* infection

## Materials and Methods

### Strains and cell culture conditions

*Francisella novicida* Utah 112 (U112) and immortalized murine bone marrow-derived macrophages were obtained from BEI Resources (NIAID, NIH, NR-13 and NR-9456) and grown according to their recommendations.

### Genetic modification of F. novicida

Genetic manipulation of *F. novicida* to generate in-frame markerless chromosomal changes was performed as previously described [3]. VSV-G (YTDIEMNRLGK) and polyhistidine (HHHHHH) fusion proteins were generated by inserting their sequences downstream of the start codon or upstream of the translation stop site. Expression of genes at a heterologous site was performed using the mini-Tn7 system as previously reported [18,14].

### Protein secretion assays

Secretion assays were performed as previously described with modifications noted below [19]. *F. novicida* cultures were diluted to OD_600_ of 0.05 in TSBC+5% KCl, grown until they reached OD_600_=1.2, and cell and supernatant fractions were separated by centrifugation. For IglC and PdpC secretion assays, 3 mL of supernatant was processed exactly as described previously [20]. For PdpE secretion, 200 mL of supernatant was filtered to remove particles greater than 0.22 µm and concentrated with a centrifugal filter with a 30kDa MWCO.

### iBMDM infection assays

Infection was performed as previously described [3]. Briefly, 4.5 × 10^4^ cells were seeded in 96-well plates in DMEM lacking antibiotics and allowed to adhere overnight. Where indicated, cells were primed with 100 ng/mL *Escherichia coli* O127:B8 lipopolysaccharide for 3 hours prior to infection. Bacteria were added at the indicated MOIs, centrifuged at 1000 G for 30 minutes, followed by a 30-minute incubation at 37ºC. After treatment with 50 µg/mL gentamycin and washing, cells were lysed with 0.1% Triton X-100 immediately and after 18 hours for CFU enumeration. For cytotoxicity measurement, lactate dehydrogenase was quantified in culture supernatants according to the manufacturer’s recommendations (Promega). IL-1β was quantified by ELISA following the manufacturer’s recommendations (BioLegend).

### Immunoprecipitation-mass spectrometry, pulldown assays, and western blots

Immunoprecipitation experiments were performed as previously reported [21]. Briefly, overnight cultures of *F. novicida* were diluted 1:100 in 1 L TSBC and grown to late log phase. Cultures were centrifuged at 6000 G for 15 minutes and pellets were resuspended in 10 mL IP buffer (20 mM Tris HCl pH 7.5, 150 mM NaCl, 1 mM AEBSF, 10 μM leupeptin, 1 μM pepstatin) supplemented with 1 mg/mL lysozyme. Bacteria were then sonicated followed by centrifugation at 4°C at 40,000 G for 30 minutes. Supernatants were incubated with 30 μL anti-VSV-G agarose or Ni-NTA beads (Sigma Aldrich) for 1 hour at 4°C, washed three times with 10 mL of IP buffer, washed further with 20 mM NH4HCO3, and then incubated 100 ng of sequencing grade trypsin for 3-6 hours. Trypsinized proteins were reduced with 1 mM TCEP, alkylated with 10 mM iodoacetamide, and purified with C18 spin columns (Pierce) prior to analysis by mass spectrometry.

Western blotting was performed using mouse α-IglC, mouse α-PdpB, rabbit α-PdpC (BEI Resources; NR-3198, NR-3196, and NR-4379, respectively), rabbit α-VSV-G (Sigma Aldrich), and mouse α-His (R&D Systems). After washing, membranes were probed with α-mouse or α-rabbit antibodies conjugated to horseradish peroxidase (Sigma Aldrich) and visualized with chemiluminescent reagent (Bio-Rad).

### *Galleria mellonella* larvae infections

Fifth instar *G. mellonella* larvae (Vanderhorst Wholesale) were sanitized with 70% ethanol and injected in the second left proleg with 10 µL containing 10^4^ CFU of log-phase bacteria suspended in PBS using a 100 µL syringe fitted with a 25G needle. After infection, larvae were singly housed in 6-well cell culture dishes at 35ºC and observed twice daily for death, defined as no movement upon stimulation.

## Acknowledgements

We wish to thank Stephanie Shames and Leah Eshraghi for critically reviewing the manuscript and Joseph D. Mougous for providing access to resources. Additionally, we wish to acknowledge the University of Florida Office of Research and College of Veterinary Medicine for funding.

